# Locus coeruleus spiking differently correlates with somatosensory cortex activity and pupil diameter

**DOI:** 10.1101/2020.06.04.134676

**Authors:** Hongdian Yang, Bilal A. Bari, Jeremiah Y. Cohen, Daniel H. O’Connor

**Affiliations:** Department of Molecular, Cell and Systems Biology, University of California, Riverside, CA 92521, USA; Department of Neuroscience, Brain Science Institute, and Kavli Neuroscience Discovery Institute, Johns Hopkins School of Medicine, Baltimore, MD, 21205, USA

## Abstract

We examined the relationships between activity in the locus coeruleus (LC), activity in the primary somatosensory cortex (S1), and pupil diameter in mice performing a tactile detection task. While LC spiking consistently preceded S1 membrane potential depolarization and pupil dilation, the correlation between S1 and pupil was more heterogeneous. Furthermore, the relationships between LC, S1 and pupil varied on timescales of sub-seconds to seconds within trials. Our data suggest that pupil diameter can be dissociated from LC spiking and cannot be used as a stationary index of LC activity.

## Intro

Multiple lines of evidence implicate the locus coeruleus/norepinephrine (LC/NE) system in perceptual task performance. First, LC activity modulates feedforward processing of sensory stimuli^1–3^, and impacts sensory cortex states^4,5^. Second, LC activity correlates with task performance^6,7^ and pupil diameter^7–10^. Finally, pupil diameter is thought to index arousal and has been found to be correlated with neuronal and behavioral detection or discrimination sensitivity^11–17^. Since sensory cortex activity impacts perceptual reports^18,19^, these observations suggest the hypothesis that LC/NE modulates sensory cortex activity and affects perceptual task performance, and that this effect can be monitored noninvasively via the easy-to-measure pupil diameter. Testing this hypothesis requires simultaneous measurement of (1) LC activity, (2) cortical activity, ideally subthreshold membrane potential, and (3) pupil diameter, all during perceptual task performance. Here, we recorded spiking activity of optogenetically-tagged LC units together with pupil diameter in mice performing a tactile detection task^20^. In a subset of experiments, we also performed simultaneous whole-cell current clamp recordings in S1 (Fig. 1).

**FIGURE 1:**
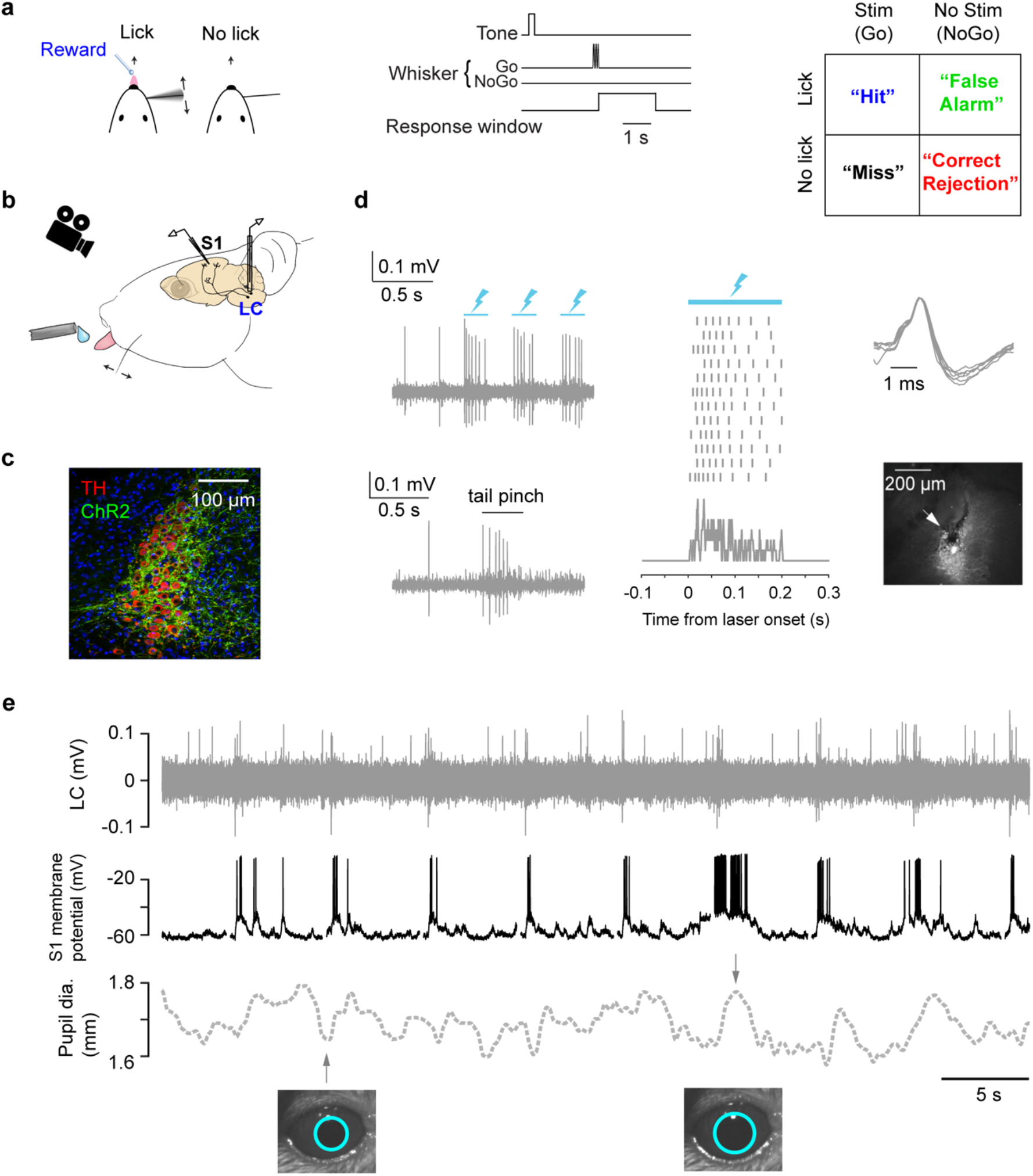
Cortical membrane potential, LC spike rate, and pupil recorded during a tactile detection task. (a) Task schematic, trial structure and all trial types of the single-whisker detection task^20^. (b) Schematic of tetrode recording in LC, whole-cell recording in S,1 and pupil tracking during the task. (c) Expression of ChR2 in a Dbh;Ai32 mouse. (ChR2-EYFP: green; Tyrosine Hydroxylase, TH: red). (d) Left: Responses of a ChR2-expressing LC unit to opto-tagging (lightning bolts: blue light pulses) and tail pinch. Middle: LC unit responses to 12 blue light pulses (200-ms) aligned to individual pulse onset. Ticks represent spikes. PSTH is shown at the bottom. Right: Typical wide waveforms of LC units and an electrolytic lesion (arrow: lesion site) in the LC (white) showing the recording location. (e) Example simultaneously recorded LC activity, S1 V_m_, and pupil.

## Results

First, we report the analysis of LC and pupil recordings during behavior (e.g., Fig. 2a). Consistent with prior reports^8–10^, cross-correlogram analysis revealed that LC spiking activity and pupil diameter were correlated across entire sessions, with pupil dilation following LC spikes (peak correlation coefficient: 0.15 ± 0.02; time lags: 2.61 ± 0.39 s, n = 39, Fig. 2b). Mean LC spiking activity aligned to trial onsets showed prominent responses to a tone delivered at the beginning of each trial, as well as in trials where mice made Go (licking) responses (Hit and False Alarm trials, Fig. 2a, c). LC spiking activity to the tone was comparable to Go responses (P = 0.24, Fig. 2d, Methods). On Hit trials, where mice successfully licked to the whisker stimulus, pre-stimulus LC activity (measured in a 0.5-s window prior to stimulus onset) was slightly but significantly lower than Miss trials, where mice failed to lick to the whisker stimulus (Fig. 2e). We note that on Miss trials LC responded weakly to whisker stimulus alone (< 0.5 sp/s above baseline, Fig. S1). LC activity measured in a short window (0.2-s) after stimulus onset was larger on Hits compared with Misses (Fig. 2e; the same trend holds for 0.1-s window, data not shown). Ideal-observer analysis showed that both pre- and post-stimulus LC activity significantly predicted perceptual reports of the mice on a trial-by-trial basis, with choice probabilities^20^ of 0.47 ± 0.014 (P = 0.032, n = 43) for pre-stimulus and 0.59 ± 0.017 (P = 4.6e-6, n = 43) for post-stimulus LC activity, respectively (Fig. 2e). LC activity aligned to the time of licking showed that spiking responses began ~200 ms prior to licking (Fig. 2f).

**FIGURE 2:**
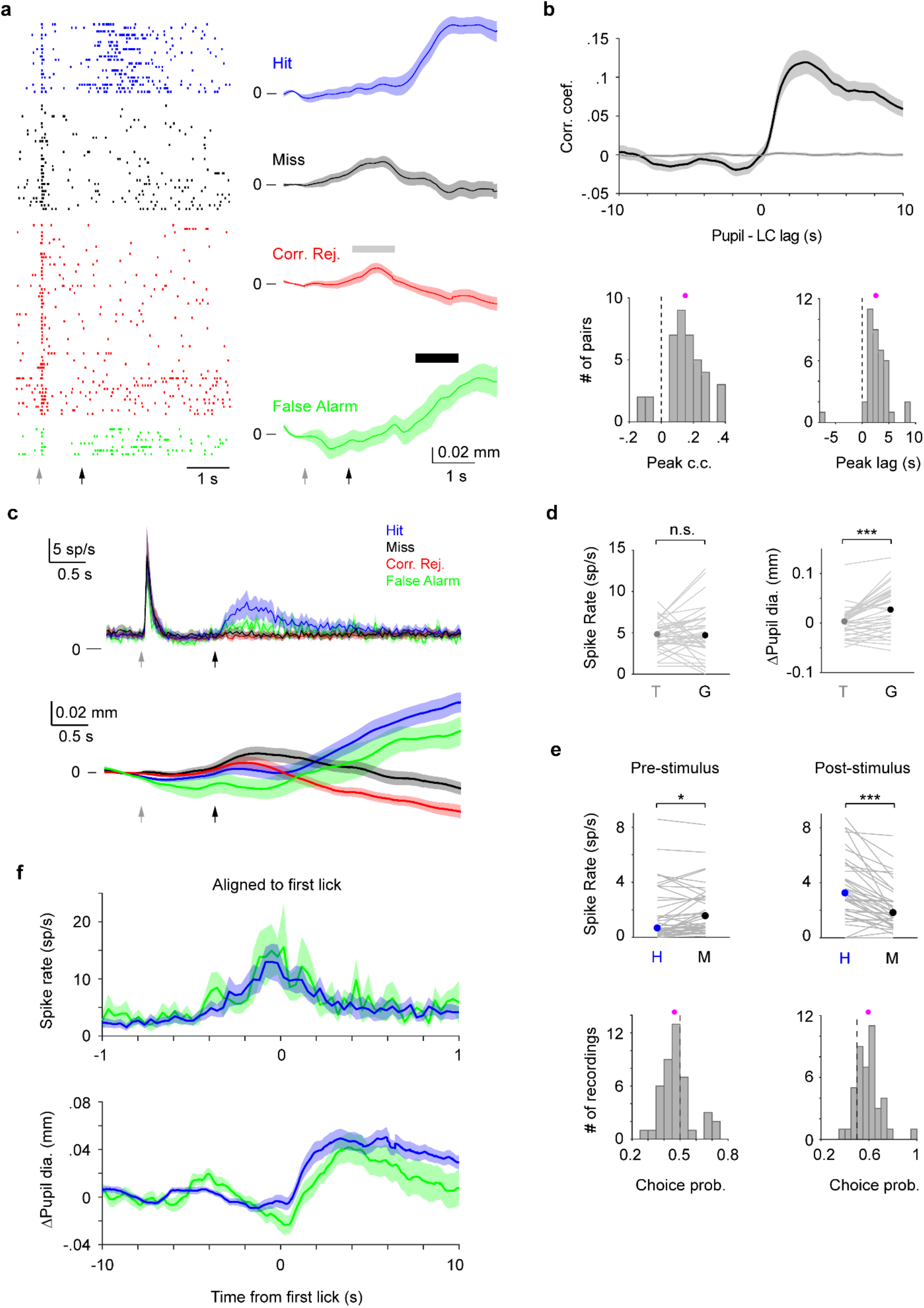
LC and pupil responses during behavior. (a) Example LC recording with pupil tracking. Left: LC spike raster separated by trial types. Right: Mean pupil diameter (± s.e.m.) separated by trial types. Grey and black arrows indicate tone and stimulus onsets, respectively. Grey and black bars indicate the time windows during which pupil responses to tone and to Go (behavioral responses) were quantified, respectively. We note that based on the temporal profiles of pupil diameter in different trial types (i.e., in the presence or absence of tactile stimulus or licking), and that tactile stimulus starts 1 s after tone onset, pupil responses to tone and Go can be segregated (Methods). (b) Top: Cross-correlogram between LC spike train and pupil diameter. Individual LC spikes were convolved with a 400-ms wide Gaussian kernel. Spike times were shuffled and LC-pupil correlations computed to establish controls (narrow grey band around zero). Bottom: Histogram of peak correlation coefficient (left), and time lags (right) between LC spike train and pupil diameter for each paired recording (magenta dot: mean). Both distributions are significantly larger than 0 (peak correlation coefficient: 0.15 ± 0.02, P = 8.3e-7, Signed rank = 743; time lags: 2.61 ± 0.39 s, P = 7.8e-7, Signed rank = 744, n = 39). (c) Trial-aligned LC spike rate (top), and pupil diameter (bottom) averaged by different trial types. Grey and black arrows indicate tone and stimulus onsets, respectively. (d) Left: LC responses to tone (T) and Go responses (G) during Hit trials with median indicated. Tone vs. Go: 4.79 (3.70 – 6.66) sp/s vs. 4.68 (3.33 – 7.26) sp/s, median (IQR), P = 0.24, Signed rank = 496.5, n = 43. Right: Pupil responses to tone and Go responses during Hit trials with median indicated. Tone vs. Go: 0.003 (−0.015 – 0.015) mm vs. 0.027 (−0.010 – 0.063) mm, median (IQR), P = 6.4e-5, Signed rank = 559, n = 36. Grey lines indicate individual recordings. (e) Top: Pre-stimulus (baseline) and post-stimulus (evoked) LC spike rate for Hit and Miss trials with median indicated (Baseline: Hit vs. Miss, 0.66 (0.30 – 3.51) sp/s vs. 1.55 (0.68 – 3.00) sp/s, median (IQR), P = 0.0083, Signed rank = 254.5; Evoked: Hit vs. Miss, 3.24 (1.78 – 5.49) sp/s vs. 1.82 (0.95 – 3.45) sp/s, median (IQR), P = 5.5e-7, Signed rank = 782.5, n = 43). Grey lines indicate individual recordings. Bottom: Histogram of choice probability for Hit vs. Miss trials based on baseline and evoked LC activity (magenta dots: mean). Choice probabilities are significantly deviated from 0.5. Baseline: 0.47 ± 0.014, P = 0.032, Signed rank = 295.5; Evoked: 0.59 ± 0.017, P = 4.6e-6, Signed rank = 751, n = 43. (f) Lick-aligned LC spike rate (top) and pupil diameter (ΔPupil, bottom) averaged by trial types: Hit (blue), FA (green).

In striking contrast, pupil diameter minimally increased in response to the tone. Instead, pupil strongly dilated on Hit and False Alarm trials, in which mice made Go (licking) responses (Fig. 2a, c, d; tone vs. Go: P = 6.4e-5, n = 36, Methods)^15^. Interestingly, pupil response to the tone was larger on Misses compared to Hits, and significantly predicted perceptual choices of the mice (Fig. S2). Pupil diameter changes (ΔPupil) aligned to the time of licking showed that pupil responses occurred after licking (Fig. 2f).

Together, these data show that LC and pupil responses were positively correlated. Both LC activity and pupil diameter increased during licking responses, but LC also strongly responded to the tone, a salient sensory cue that alerted mice of trial onsets. Thus, LC activity and pupil diameter appear to reflect different sets of task events during this behavior.

Next, we analyzed recordings where we simultaneously measured membrane potential (V_m_) of S1 neurons (mostly from layer 2/3, Fig. S3) along with LC spiking and/or pupil diameter during the detection task. Our goal was to determine how LC spiking related to cortical activity and to pupil diameter during task performance. We used spike-triggered averages (STAs) to quantify how individual spikes from single LC units correlated with changes in V_m_ and pupil diameter. LC spike-triggered V_m_ analyses revealed that LC spikes were associated with a depolarization in cortical neurons (1.39 ± 0.35 mV, n = 12, Fig. 3a-c). On average, V_m_ depolarization associated with an LC spike peaked after the spike, with short time lags from an LC spike to peak depolarization in S1 (0.17 ± 0.06 s, n = 12, Fig. 3a-c, also see Fig. S4).

**FIGURE 3:**
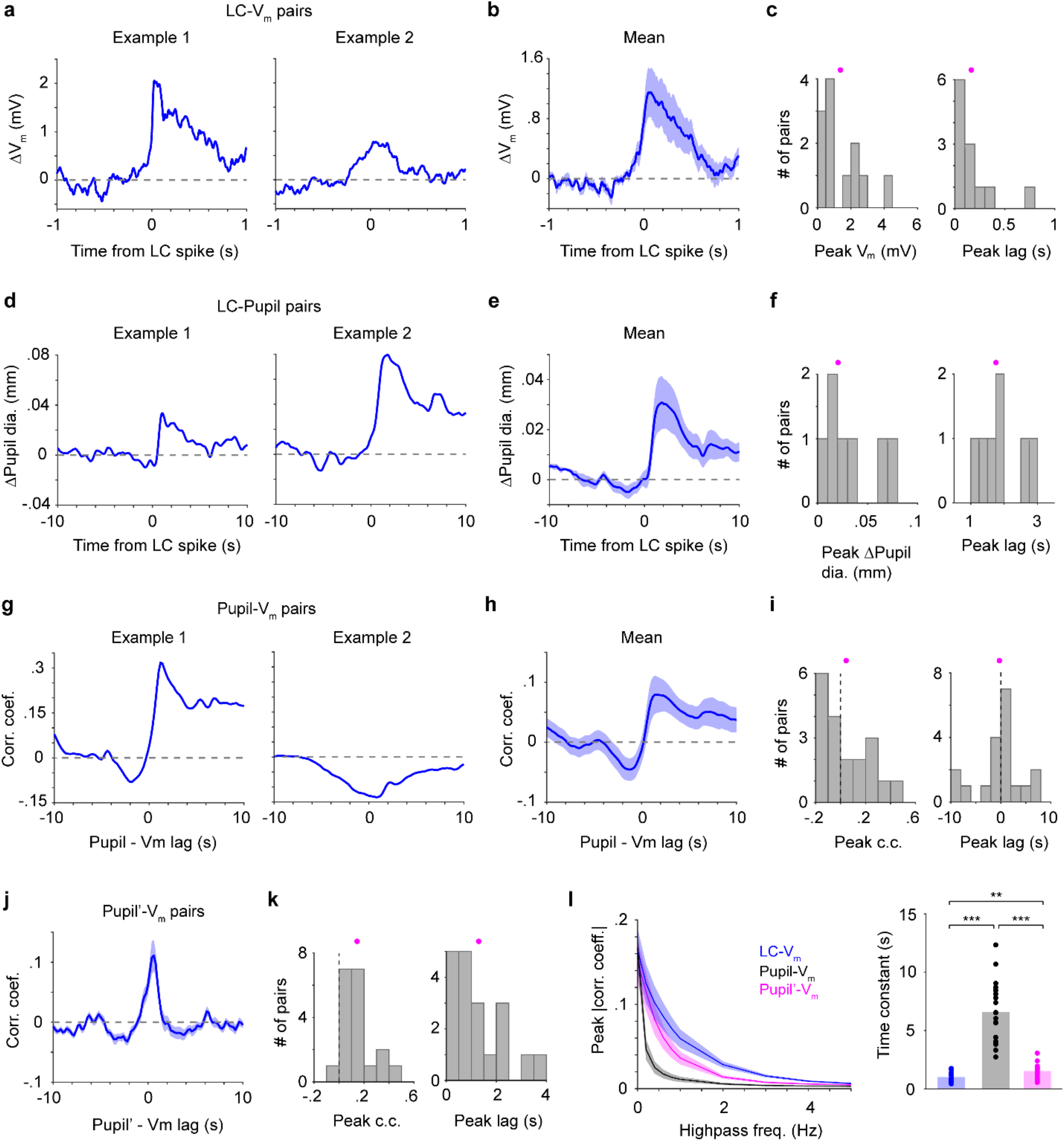
Different relationships between LC spikes, S1 V_m_ and pupil diameter. (a) Two examples of LC spike-triggered average ΔV_m_. (b) Group mean of LC spike-triggered average ΔV_m_ (± s.e.m., n = 12) (c) Histograms of peak ΔV_m_ and peak lags (showing all LC-S1 pairs) with means indicated (magenta dots). Both distributions are significantly larger than 0 (Peak ΔV_m_: 1.39 ± 0.35 mV, P = 4.9e-4, Signed rank = 78; Peak lags: 0.17 ± 0.06 s, P = 4.9e-4, Signed rank = 78, n = 12). (d) Two examples of LC spike-triggered average ΔPupil. (e) LC spike-triggered average ΔPupil group mean (± s.e.m., n = 7). (f) Histograms of peak ΔPupil and peak lags (showing all LC-Pupil pairs) with means indicated (magenta dots). Both distributions are significantly larger than 0 (Peak ΔPupil: 0.03 ± 0.01 mm, P = 0.016, Signed rank = 28; Peak lags: 1.89 ± 0.25 s, P = 0.016, Signed rank = 28, n = 7). (g) Two examples of Pupil-V_m_ cross-correlograms. (h) Group mean of Pupil-V_m_ cross-correlograms (± s.e.m., n = 19). (i) Histograms of peak Pupil-V_m_ correlation coefficient and peak lags (showing all S1-Pupil pairs) with means indicated (magenta dots). Both distributions are not significantly deviated from 0 (Peak correlation coefficient: 0.05 ± 0.04, P = 0.33, Signed rank = 119; Peak lags: - 0.22 ± 1.01 s, P = 0.87, Signed rank = 99, n = 19). (j) Group mean of the time derivative of pupil (Pupil’)-V_m_ cross-correlograms (± s.e.m., n = 19). (k) Histograms of peak Pupil’-V_m_ correlation coefficient and peak lags with means indicated (magenta dots). Both distributions are significantly larger than 0 (Peak correlation coefficient: 0.15 ± 0.03, P = 1.6e-4, Signed rank = 189; Peak lags: 1.31 ± 0.24 s, P = 1.3e-4, Signed rank = 190, n = 19). (l) Left: Peak correlation coefficient for LC-V_m_, Pupil-V_m_ and Pupil’-V_m_ pairs after progressive high-pass filtering of S1 V_m_. Right: Exponential decay functions (corr. coef. = a*exp(−freq*μ)) were fitted to these curves. The time constant μ is significantly different (repeated-measures ANOVA, F(2, 36) = 74.5, P = 1.6e-13, n = 19). Post-hoc Tukey-Kramer tests revealed that the LC-V_m_ relationship had the slowest decay and Pupil-V_m_ had the fastest decay. LC-V_m_ vs. Pupil-V_m_, P = 5.9e-8; LC-V_m_ vs. Pupil’-V_m_, P = 0.0037; Pupil-V_m_ vs. Pupil’-V_m_, P = 7.1e-7.

Consistent with the previous cross-correlogram analysis based on a larger set of LC-pupil recordings (Fig. 2b), here STA analysis showed that pupil diameter increased in association with individual spikes from LC single units (0.03 ± 0.01 mm, n = 7), with peak dilation occurring roughly ten-fold slower than peak V_m_ depolarization (time lags from an LC spike to peak pupil dilation: 1.89 ± 0.25 s, n = 7, Fig. 3d-f).

Given that pupil diameter and LC activity are positively correlated, and that pupil diameter has been often considered to index LC activity^16,21^, we next tested whether the pupil-S1 relationship resembled the LC-S1 relationship. Cross-correlogram analyses revealed heterogeneous correlations between pupil diameter and S1 V_m_, with both positive and negative correlations as well as positive and negative time lags (peak correlation coefficient: 0.05 ± 0.04; time lags: - 0.22 ± 1.01 s, n = 19, Fig. 3g-i). In sharp contrast, the time derivative of pupil diameter (pupil’) was positively correlated with S1 V_m_ in a more consistent manner (peak correlation coefficient: 0.15 ± 0.03; time lags: 1.31 ± 0.24 s, n = 19, Fig. 3j,k)^10,12^. We further examined how well LC spiking and pupil diameter can predict cortical V_m_ fluctuations at different timescales. We found that LC activity was superior in predicting cortical dynamics faster than ~200-300 ms (exponential decay time constant: LC-V_m_ vs. Pupil-V_m_ vs. Pupil’-V_m_,1.02 ± 0.09 vs. 6.59 ± 0.60 vs. 1.53 ± 0.14, Fig. 3l; repeated-measures ANOVA, F(2, 36) = 74.5, P = 1.6e-13, n = 19). Post-hoc Tukey-Kramer tests revealed that the LC-V_m_ relationship had the slowest decay and Pupil-V_m_ had the fastest decay (LC-V_m_ vs. Pupil-V_m_, P = 5.9e-8; LC-V_m_ vs. Pupil’-V_m_, P = 0.0037; Pupil-V_m_ vs. Pupil’-V_m_, P = 7.1e-7). On the other hand, the LC-Pupil’ relationship was very similar to that of LC-Pupil (compare Fig. S5a,b with Fig. 3e,f).

Together, these data show that LC spikes preceded S1 depolarizations and pupil dilations. LC spiking correlated with both V_m_ and pupil diameter changes, but on vastly different timescales (~0.2 s vs. ~2 s). Our data also show that the time derivative of pupil diameter, but not the absolute pupil size, is a good predictor of S1 V_m_ fluctuations. However, LC spiking can track fast V_m_ fluctuations better than either pupil and pupil’.

Individual trials in our detection task contained distinct events, including the tone that alerted mice of the trial start (“Tone”), the whisker stimulus on Go trials (“Stimulus”), and licks (“Lick”), as well as other periods in which mice did not receive stimuli or make lick responses (“Quiet”). For a more granular perspective on how LC spiking correlated with changes in V_m_ and pupil diameter, we computed LC spike-triggered averages separately in these different event windows (task epochs, Methods).

While single LC spikes were associated with prominent changes in both cortical V_m_ and pupil diameter, we found that these associations strikingly depended on task epochs: V_m_ depolarization associated with an LC spike had the biggest response to tone/licking and almost no response during the quiet periods (Fig. 4a). In contrast, pupil dilation associated with an LC spike had the biggest response to licking and almost no response to the tone (Fig. 4b). The pupil’ associated with an LC spike had an intermediate response to the tone (Fig. 4c). In addition, peak pupil dilation, pupil’ and V_m_ depolarization appeared to have different dependencies on LC spike counts, with a roughly monotonic relationship between pupil and LC, and a much weaker dependence between V_m_ and LC (Fig. S6). Thus, the correlations between LC spiking and V_m_, and between LC spiking and pupil diameter, are non-stationary, even on the timescale of a few seconds. Importantly, these epoch-dependencies were different for V_m_ and pupil - with the biggest response occurring to the tone for V_m_, and the smallest response occurring to the tone for pupil - suggesting that the correlations between LC activity and V_m_ and pupil each reflect distinct unmeasured underlying processes.

**FIGURE 4:**
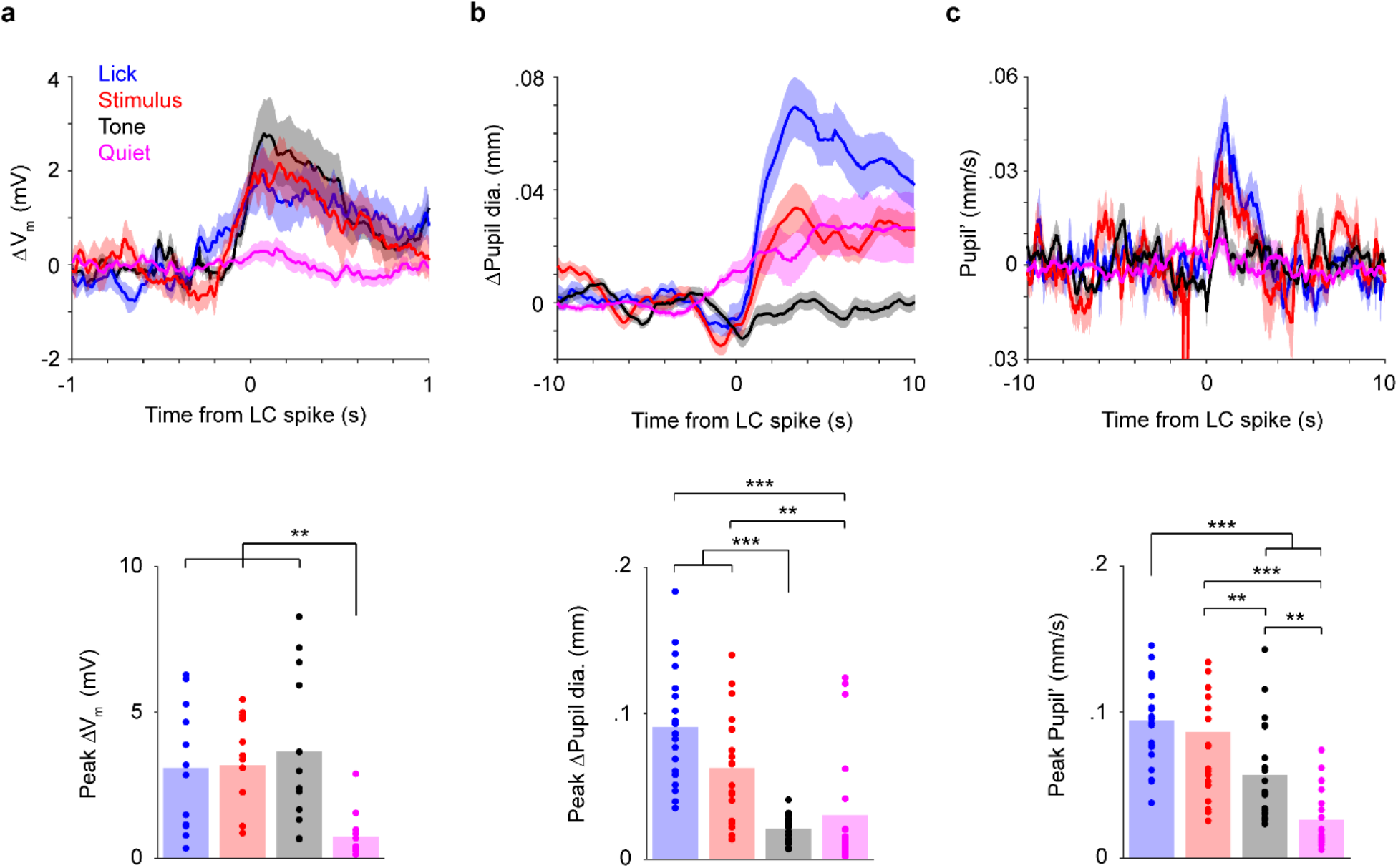
Correlations between LC spikes, S1 V_m_ and pupil diameter depend on task epoch. (a) Top: LC spike-triggered ΔV_m_ separated by task epoch: tone, stimulus, lick and quiet. Bottom: Bar graphs of peak ΔV_m_ for each epoch. Dots indicate individual paired recordings. Repeated-measure ANOVA, F(3, 33) = 9.2, P = 1.4e-4, n = 12. Post-hoc Tukey-Kramer tests revealed that peak ΔV_m_ in lick, stimulus and tone epochs were not different from each other. Lick vs. Stim, P = 1.00; Lick vs. Tone, P = 0.76; Stim vs. Tone, P = 0.94. Peak ΔV_m_ in quiet epochs was lower. Quiet vs. Lick, P = 0.0059; Quiet vs. Stim, P = 0.0038; Quiet vs. Tone, P = 0.0041. (b) Top: LC spike-triggered ΔPupil separated by task epoch. Bottom: Bar graphs of peak ΔPupil for each epoch. Dots indicate individual paired recordings. Repeated-measure ANOVA, F(3, 57) = 22.1, P = 1.3e-9, n = 20. Post-hoc Tukey-Kramer tests revealed that peak ΔPupil in lick and stimulus epochs were larger than in tone and quiet epochs. Lick vs. Stim, P = 0.10; Tone vs. Quiet, P = 0.76; Lick vs. Tone, P = 3.7e-7; Lick vs. Quiet, P = 6.2e-4; Stim vs. Tone, P = 1.1e-4; Stim vs. Quiet, P = 0.0027. (c) Top: LC spike-triggered pupil’ separated by task epoch. Bottom: Bar graphs of peak pupil’ for each epoch. Dots indicate individual paired recordings. Repeated-measures ANOVA, F(3, 57) = 35.3, P = 4.9e-13, n = 20. Post-hoc Tukey-Kramer tests revealed that peak pupil’ in lick and stimulus epochs were larger than in tone, and peak pupil’ in quiet epochs was the lowest. Lick vs. Stim, P = 0.46; Tone vs. Quiet, P = 0.0013; Lick vs. Tone, P = 1.0e-4; Lick vs. Quiet, P = 1.4e-8; Stim vs. Tone, P = 0.0058; Stim vs. Quiet, P = 6.7e-6.

## Discussion

We found that pre-stimulus baseline LC spiking predicted behavioral responses. Thus, fluctuations in LC/NE activity may in part underlie perceptual task performance. However, the effect was weak, possibly due to the use of an auditory cue that puts the mice in a more homogeneous arousal state. As a result, factors other than fluctuations of arousal also likely contribute to cortical choice probabilities observed in prior work with this task^20^. In other tasks without such alerting cues, task performance may have a stronger dependence on arousal and pre-stimulus LC activity.

LC responded strongly to an auditory cue (tone) meant to alert the mice to the beginning of a trial. While this tone carried no information about the presence of a tactile stimulus or reward on any given trial, and therefore was not associated with a particular movement response, it did inform the mice about the time when a tactile stimulus could occur (in our task the duration between the tone and stimulus onset was fixed). The robust LC spiking responses to this cue are therefore consistent with LC’s role in promoting alertness or preparedness to detect a weak stimulus. We also found that LC responded to operant licking responses, which is consistent with earlier work showing that LC encoded overt decision execution^22^.

Our data show that while LC spiking and pupil diameter correlate well at long timescales, and both can predict changes in cortical dynamics, LC does so an order of magnitude faster. Moreover, the correlation between pupil and V_m_ is much more heterogeneous than between LC and V_m_. In support of previous studies, our results suggest that compared with change in the absolute size of pupil diameter, its time derivative is a better predictor of cortical states^10,12^. Importantly, the relationships between LC activity, S1 V_m_ and pupil depended on task epoch. Because these epochs changed on the timescale of a few seconds, our data imply that pupil diameter can be dissociated from LC spiking and cannot be used as a stationary index of LC activity. However, comparing across repeats of similar epochs should yield a more accurate prediction of LC spiking by pupil diameter. That is, in attempting to use pupil diameter as a proxy for LC spiking, our data suggest it would be useful to separately normalize distinct task epochs. Future work should examine the LC-pupil relationship using fine-scale analyses that consider the behavioral states at a granular level specific to individual tasks.

## Author Contributions

H.Y. performed all experiments with help from B.A.B. H.Y., B.A.B. and D.H.O. analyzed data. H.Y., J.Y.C. and D.H.O. planned the project. H.Y. and D.H.O. wrote the paper with input from B.A.B. and J.Y.C.

## Acknowledgments

We thank E. Zagha for comments on the manuscript; Dwight E. Bergles for DBH-Cre mice. This work was supported by UCR startup (H.Y.), Klingenstein-Simons Fellowship Awards in Neuroscience (H.Y.), NIH grants F30MH110084 (B.A.B.), 1R01NS107355 (H.Y.), 1R01NS112200 (H.Y.), R01NS089652 (D.H.O.), 1R01NS104834-01 (D.H.O., J.Y.C), and P30NS050274.

## Methods

All procedures were performed in accordance with protocols approved by the Johns Hopkins University Animal Care and Use Committee. Mice were DBH-Cre (B6.FVB(Cg)-Tg(Dbh-cre) KH212Gsat/Mmucd, 036778-UCD, MMRRC); Ai32 (RCL-ChR2(H134R)/EYFP, 012569, JAX), singly housed in a vivarium with reverse light-dark cycle (12 hr each phase). Male and female mice of 6-12 weeks were implanted with titanium head posts as described previously^20^. After recovery, mice were trained to perform a Go/NoGo single whisker detection task as described previously^20^.

Custom microdrives with eight tetrodes and an optic fiber^23^ (0.39 NA, 200 um core) were built to make extracellular recordings from LC neurons. Each tetrode comprised four nichrome wires (100-300 KΩ). A ~1 mm diameter craniotomy was made (centered at −5.2 mm caudal and 0.85 mm lateral relative to bregma) for implanting the tetrodes to a depth of 2.7 mm relative to the brain surface. The microdrive was advanced in steps of ~100 um each day until reaching LC, identified by optogenetic tagging of DBH+ neurons expressing ChR2, tail pinch response, wide extracellular spike waveforms and post-hoc electrolytic lesions. Broadband voltage traces were acquired at 30 kHz (Intan Technologies) and filtered between 0.1 and 10 kHz. Signals were then bandpass filtered between 300 and 6000 Hz, and spikes were detected using a threshold of 4-6 standard deviations. The timestamp of the peak of each detected spike, as well as a 1-ms waveform centered at the peak were extracted from each channel for offline spike sorting using MClust^24^. At the conclusion of the experiments, brains were perfused with PBS followed by 4% PFA, post-fixed overnight, then cut into 100 μm coronal sections and stained with anti-Tyrosine Hydroxylase (TH) antibody (Millipore AB152).

Pupil video was acquired at 50 Hz using a PhotonFocus camera and StreamPix 5 software. Light from a 940 LED was passed through a condenser lens and directed to the right eye, reflected off a mirror, and directed into a 0.25X telecentric lens. WaveSurfer (https://www.janelia.org/open-science/wavesurfer) triggered individual camera frames synchronized with electrophysiological recordings.

In a subset of animals, we performed simultaneous intracellular current clamp (whole-cell) recordings in conjunction with LC recording and/or pupil tracking during behavior. A craniotomy over the C2 barrel was made based on intrinsic signal imaging^20^. In some cases, we also made craniotomies over nearby barrels based on the known somatotopy of S1^25,26^ to increase yield. Whole-cell recording procedures, quality control and data processing were performed as described previously^20^.

For Fig. 2d, LC responses to the tone were calculated using a 300-ms window starting at tone onset, and LC responses to Go were calculated using a 300-ms window starting 200 ms after stimulus onset to capture peak responses. These estimates were based on LC response profiles in Fig. 2c. Pupil responses to the tone were calculated using a 1-s window starting 1 s after tone onset. This estimate was primarily based on pupil response profile during CR trials (e.g., Fig. 2a, c, indicated by the grey bar), where there was no whisker stimulus or licking response. Pupil responses to Go (licking) were calculated using a 1-s window starting 1.5 s after stimulus onset (e.g., FA trials in Fig. 2a, c, indicated by the black bar). Based on the temporal profiles of pupil diameter in different trial types shown in Fig. 2a, c, and that the whisker stimulus started 1 s after tone onset, pupil responses to tone and Go can be segregated. These estimates were consistent with the results showing that pupil dilated 1-2 s after LC spikes (Fig. 2b, and Fig. 3d-f).

For Fig. 2e, pre-stimulus LC baseline activity was calculated using a 500-ms window ending 50 ms before stimulus onset. Post-stimulus activity was calculated using a 200-ms window starting 20 ms after stimulus onset, before licking responses^20^. Choice probabilities were computed as described previously^20^.

To compute lick-aligned changes in LC spiking and pupil diameter, we only used licks that occurred at least 0.5 s after the previous lick. To compute LC spike triggered S1 V_m_ and pupil, we only used LC spikes that occurred at least 0.5 s after the previous spike. For STA analysis, peak ΔV_m_, ΔPupil or the time derivative of Pupil (Pupil’) was defined as the largest positive or negative value within the observed window (± 1 s or ± 10 s, respectively).

For cross-correlogram analysis, each LC spike train was convolved with a 400-ms wide Gaussian kernel (results hold for 200-ms kernel, data not shown). Peak correlation coefficients were defined as the largest positive or negative value within the observed window (± 1 s or ± 10 s). To examine how well LC spiking and pupil diameter could predict cortical V_m_ fluctuations at different timescales (Fig. 3l), V_m_ was high-pass filtered at 0, 0.2, 0.4, 0.6, 0.8, 1, 2, 3, 4 and 5 Hz separately. Cross-correlogram analysis between the filtered V_m_ and LC (pupil) activity was then performed as described above, and largest absolute values of peak correlation coefficients were taken.

Task epochs were defined as: “Tone” epochs: −0.2 s to 0.3 s with respect to tone onset; “Stimulus” epochs: −0.2 s to 0.3 s with respect to stimulus onset (i.e., only on trials with whisker stimulation); “Licking” epochs: −0.2 s to 0.3 s with respect to licks that occurred at least 0.5 s after the previous lick; “Quiet” epochs: non-overlapping 0.5 s segments excluding the three types of epoch defined previously during the entire session.

Thirty-nine LC-pupil pairs were included in Fig. 2b, including single- and multi-units, with and without S1 recordings. For the rest of Fig. 2, LC analysis included forty-three recordings, each with at least 4 Hit and 4 Miss trials. Among those, thirty-six were with pupil recordings, and were used for pupil analysis. Twelve pairs of S1 whole-cell and LC single-unit recordings were included in Fig. 3a-c, 4a, seven of which were with pupil recordings and included in Fig. 3d-f. Nineteen S1-pupil recordings were included in Fig. 3g-l. Twenty pairs of LC SU and pupil recordings were included in Fig. 4b,c, with and without S1 recordings.

Data were reported as mean ± s.e.m. unless otherwise noted. Statistical tests were by two-tailed Wilcoxon signed rank unless otherwise noted. We did not use statistical methods to predetermine sample sizes. Sample sizes are similar to those reported in the field. We assigned mice to experimental groups arbitrarily, without randomization or blinding.

### Data availability

Data are available from the corresponding author upon request

### Code availability

MATLAB scripts used to analyze the data are available from the corresponding author upon request.

---

**Figure S1.**
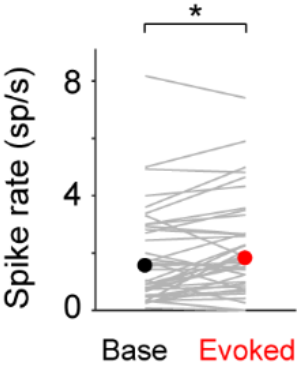
LC responded minimally to whisker stimulation when mice did not make a licking response (Miss trials, Baseline vs. Evoked: 1.55 (0.68-3.00) sp/s vs. 1.82 (0.95-3.45) sp/s, median (IQR), P = 0.02, n = 43).

**Figure S2.**
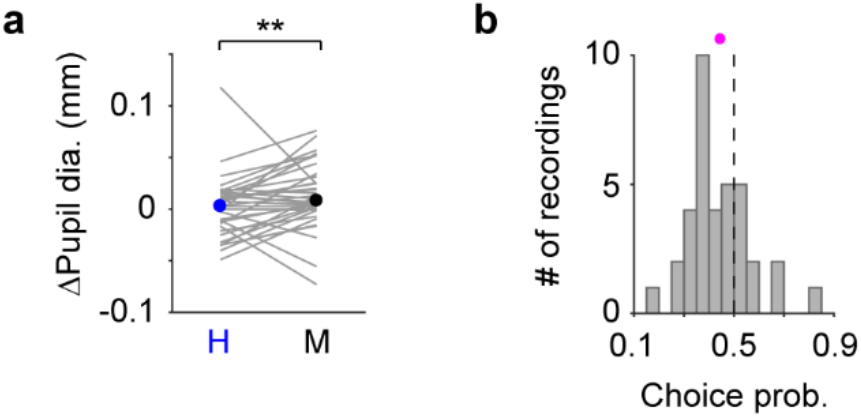
(a) Pupil responses to the tone for Hit and Miss trials with median indicated. Hit vs. Miss, 0.003 (−0.015 – 0.015) mm vs. 0.0083 (−0.0005 – 0.029) mm, median (IQR), P = 0.0062, n = 36. Grey lines indicate individual recordings. (b) Histogram of choice probability for Hit vs. Miss trials based on pupil responses to the tone (magenta dot: mean). Choice probability is significantly deviated from 0.5 (0.44 ± 0.021, P = 0.0036, n = 36).

**Figure S3.**
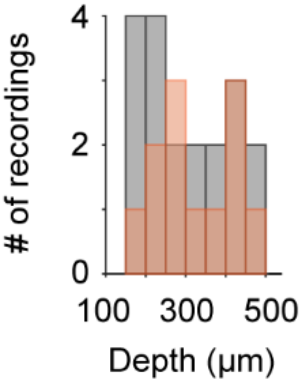
Histograms of the depth of S1 whole-cell recordings. Red: 12 S1 recordings included in the LC-S1 pairs in Fig. 3a-c, 3l, 4a. Grey: 19 S1 recordings included in the Pupil-S1 pairs in Fig. 3g-I.

**Figure S4.**
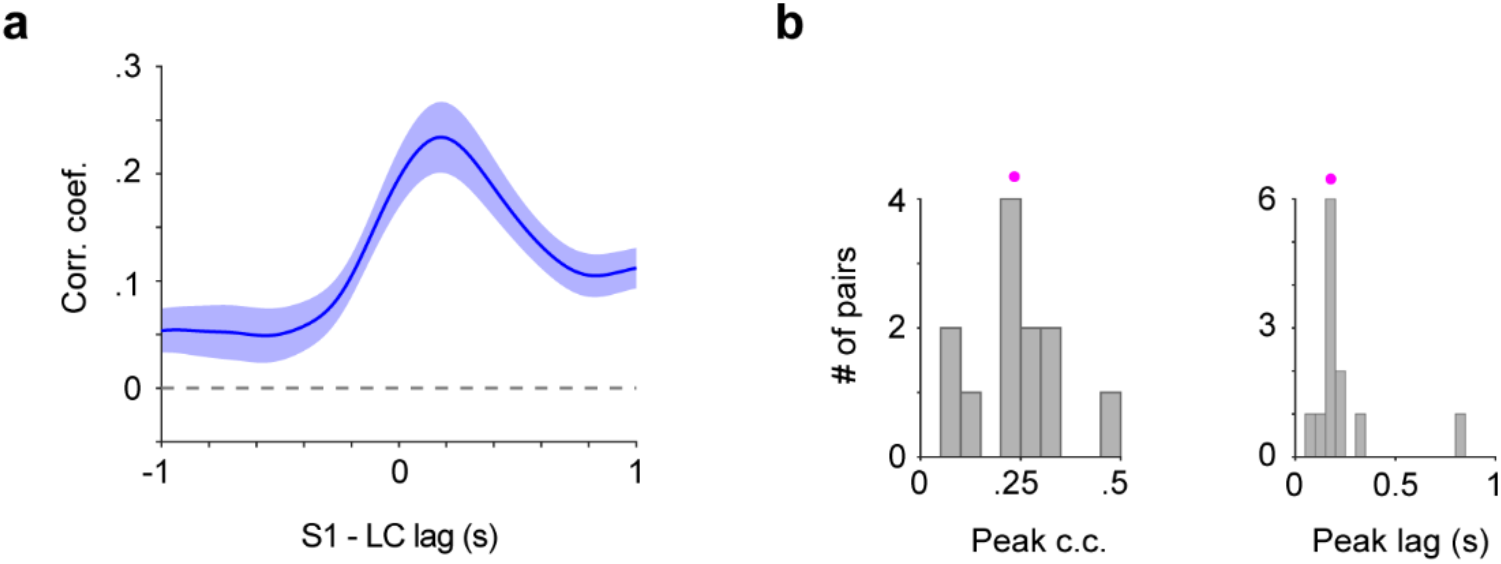
(a) Group mean of cross-correlogram between LC spike train and S1 V_m_ (n = 12). Individual LC spikes were convolved with a 400-ms wide Gaussian kernel. (b) Histogram of peak correlation coefficient (0.24 ± 0.03) and time lag (0.24 ± 0.06 s) between LC spike train and S1 V_m_ for each paired recording (pupil dot: mean).

**Figure S5.**
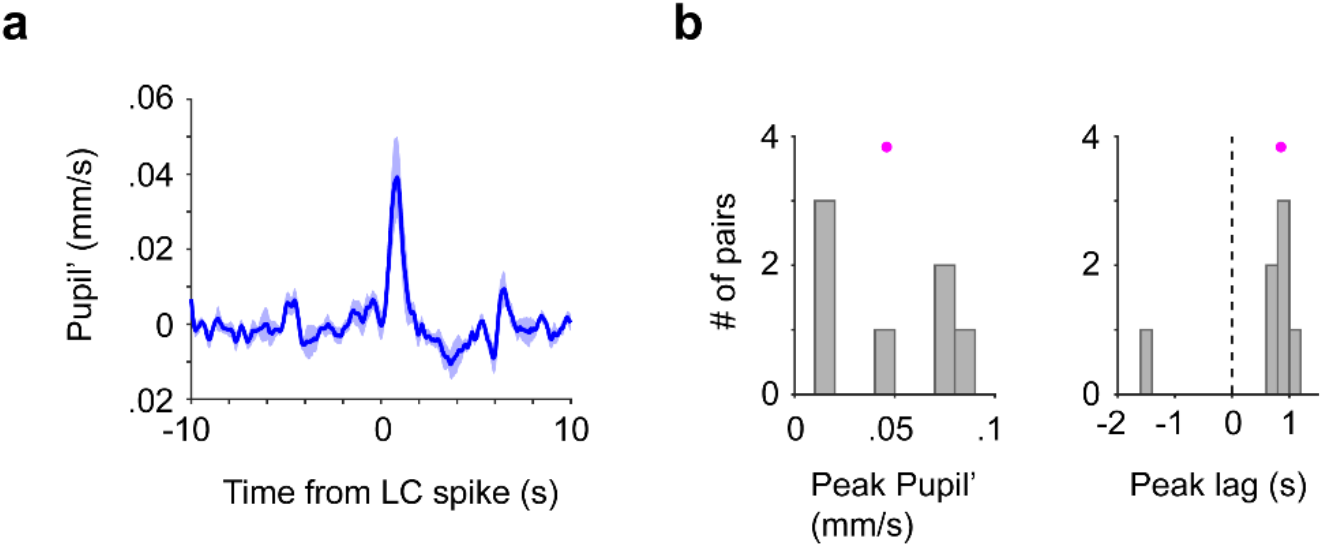
(a) Group mean of LC spike-triggered time derivative of pupil diameter (Pupil’, n = 7), same dataset as used in Fig. 3d-f. (b) Histogram of peak pupil’ (0.05 ± 0.01 mm/s) and time lag (0.53 ± 0.35 s) between LC spike train and pupil’ for each paired recording (pupil dot: mean).

**Figure S6.**
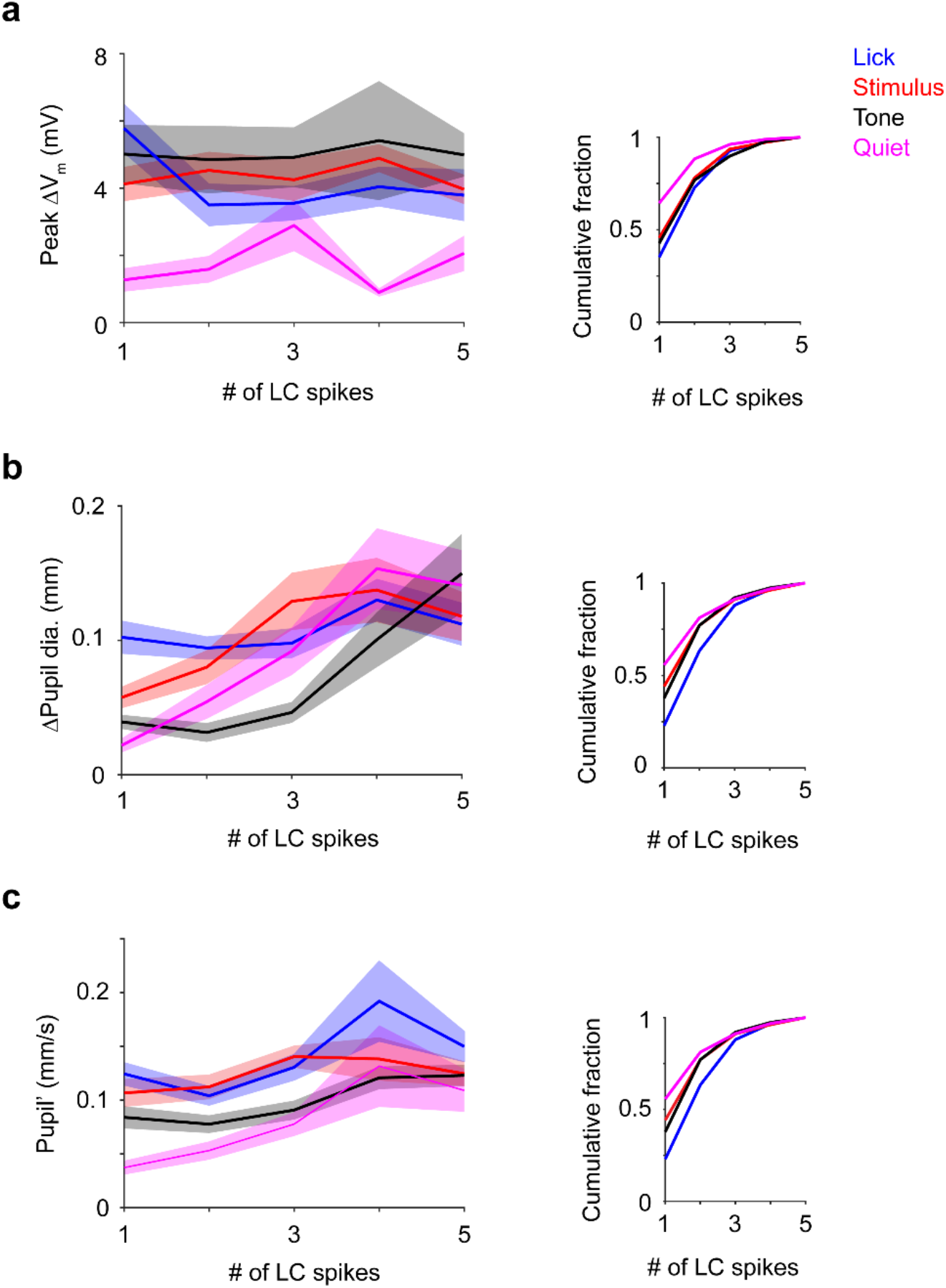
(a) Left: Peak ΔV_m_ vs. LC spike counts by epochs. Right: Cumulative histograms showing numbers of trials that go into the plots when broken down by LC spike counts. (b) Left: Peak ΔPupil vs. LC spike counts by epochs. Right: Cumulative histograms showing numbers of trials that go into the plots when broken down by LC spike counts. (c) Left: Peak Pupil’ vs. LC spike counts by epochs. Right: Cumulative histograms showing numbers of trials that go into the plots when broken down by LC spike counts, which is identical to the histogram in b.

## References

1. Rodenkirch, C., Liu, Y., Schriver, B. J. & Wang, Q. Locus coeruleus activation enhances thalamic feature selectivity via norepinephrine regulation of intrathalamic circuit dynamics. Nat. Neurosci. 22, 120–133 (2019).

2. Hirata, A., Aguilar, J. & Castro-Alamancos, M. A. Noradrenergic activation amplifies bottom-up and top-down signal-to-noise ratios in sensory thalamus. J. Neurosci. 26, 4426–36 (2006).

3. Devilbiss, D. M., Page, M. E. & Waterhouse, B. D. Locus ceruleus regulates sensory encoding by neurons and networks in waking animals. J. Neurosci. 26, 9860–72 (2006).

4. Polack, P.-O., Friedman, J. & Golshani, P. Cellular mechanisms of brain state–dependent gain modulation in visual cortex. Nat. Neurosci. 16, 1331–1339 (2013).

5. Constantinople, C. M. & Bruno, R. M. Effects and mechanisms of wakefulness on local cortical networks. Neuron 69, 1061–8 (2011).

6. Usher, M., Cohen, J. D., Servan-Schreiber, D., Rajkowski, J. & Aston-Jones, G. The role of locus coeruleus in the regulation of cognitive performance. Science 283, 549–54 (1999).

7. Rajkowski, J., Kubiak, P. & Aston-Jones, G. Locus coeruleus activity in monkey: Phasic and tonic changes are associated with altered vigilance. Brain Res. Bull. 35, 607–616 (1994).

8. Joshi, S., Li, Y., Kalwani, R. M. & Gold, J. I. Relationships between Pupil Diameter and Neuronal Activity in the Locus Coeruleus, Colliculi, and Cingulate Cortex. Neuron 89, 221–234 (2016).

9. Liu, Y., Rodenkirch, C., Moskowitz, N., Schriver, B. & Wang, Q. Dynamic Lateralization of Pupil Dilation Evoked by Locus Coeruleus Activation Results from Sympathetic, Not Parasympathetic, Contributions. Cell Rep. 20, 3099–3112 (2017).

10. Reimer, J. et al. Pupil fluctuations track rapid changes in adrenergic and cholinergic activity in cortex. Nat. Commun. 7, 13289 (2016).

11. McGinley, M. J., David, S. V. & McCormick, D. A. Cortical Membrane Potential Signature of Optimal States for Sensory Signal Detection. Neuron 87, 179–192 (2015).

12. Reimer, J. et al. Pupil Fluctuations Track Fast Switching of Cortical States during Quiet Wakefulness. Neuron 84, 355–362 (2014).

13. Vinck, M., Batista-Brito, R., Knoblich, U. & Cardin, J. A. Arousal and Locomotion Make Distinct Contributions to Cortical Activity Patterns and Visual Encoding. Neuron 86, 740–754 (2015).

14. Schriver, B. J., Bagdasarov, S. & Wang, Q. Pupil-linked arousal modulates behavior in rats performing a whisker deflection direction discrimination task. J. Neurophysiol. 120, 1655–1670 (2018).

15. Lee, C. R. & Margolis, D. J. Pupil Dynamics Reflect Behavioral Choice and Learning in a Go/NoGo Tactile Decision-Making Task in Mice. Front. Behav. Neurosci. 10, 1–14 (2016).

16. McGinley, M. J. et al. Waking State: Rapid Variations Modulate Neural and Behavioral Responses. Neuron 87, 1143–1161 (2015).

17. Lee, C. C. Y., Kheradpezhouh, E., Diamond, M. E. & Arabzadeh, E. State-Dependent Changes in Perception and Coding in the Mouse Somatosensory Cortex. Cell Rep. 32, 108197 (2020).

18. Sachidhanandam, S., Sreenivasan, V., Kyriakatos, A., Kremer, Y. & Petersen, C. C. H. Membrane potential correlates of sensory perception in mouse barrel cortex. Nat. Neurosci. 16, 1671–1677 (2013).

19. Miyashita, T. & Feldman, D. E. Behavioral detection of passive whisker stimuli requires somatosensory cortex. Cereb. Cortex 23, 1655–1662 (2013).

20. Yang, H., Kwon, S. E., Severson, K. S. & O’Connor, D. H. Origins of choice-related activity in mouse somatosensory cortex. Nat. Neurosci. 19, 127–134 (2016).

21. Aston-Jones, G. & Cohen, J. D. An integrative theory of locus coeruleus-norepinephrine function: adaptive gain and optimal performance. Annu. Rev. Neurosci. 28, 403–50 (2005).

22. Kalwani, R. M., Joshi, S. & Gold, J. I. Phasic Activation of Individual Neurons in the Locus Ceruleus/Subceruleus Complex of Monkeys Reflects Rewarded Decisions to Go But Not Stop. J. Neurosci. 34, 13656–13669 (2014).

23. Cohen, J. Y., Haesler, S., Vong, L., Lowell, B. B. & Uchida, N. Neuron-type-specific signals for reward and punishment in the ventral tegmental area. Nature 482, 85–8 (2012).

24. Redish, A. D. MClust Spike sorting toolbox Documentation for version 4.4. http://redishlab.neuroscience.umn.edu/MClust/MClust-4.4.pdf (2014).

25. Welker, C. & Woolsey, T. A. Structure of layer IV in the somatosensory neocortex of the rat: Description and comparison with the mouse. J. Comp. Neurol. 158, 437–453 (1974).

26. Wilson, M. a, Johnston, M. V, Goldstein, G. W. & Blue, M. E. Neonatal lead exposure impairs development of rodent barrel field cortex. Proc. Natl. Acad. Sci. U. S. A. 97, 5540–5545 (2000).

